# The Role of Genetic Variation in Maize Response to Beneficial Endophytes

**DOI:** 10.1101/2021.11.03.467096

**Authors:** Corey Schultz, Kamaya Brantley, Jason Wallace

## Abstract

**Background:** Growth-promoting endophytes have great potential to boost crop production and sustainability. There is, however, a lack of research on how differences in the plant host affect an endophyte’s ability to promote growth. We set out to quantify how different maize genotypes respond to specific growth-promoting endophytes. We inoculated genetically diverse maize lines with three different known beneficial endophytes: *Herbaspirillum seropedicae* (a gram-negative bacteria), *Burkholderia* WP9 (a gram-negative bacteria), and *Serendipita vermifera* Subsp. *bescii* (a Basidiomycota fungus). Maize seedlings were grown for 3 weeks under controlled conditions in the greenhouse and assessed for various growth promotion phenotypes.

**Results:** We found *Herbaspirillum seropedicae* to increase chlorophyll content, plant height, root length, and root volume significantly in different maize genotypes, while *Burkholderia* WP9 did not significantly promote growth in any lines under these conditions. *Serendipita bescii* significantly increased root and shoot mass for 4 maize genotypes, and growth promotion correlated with measured fungal abundance. Although plant genetic variation by itself had a strong effect on phenotype, its interaction with the different endophytes was weak, and the endophytes rarely produced consistent effects across different genotypes.

**Conclusions:** This genome-by-genome interaction indicates that the relationship between a plant host and beneficial endophytes is complex, and it may partly explain why many microbe-based growth stimulants fail to translate from laboratory settings to the field. Detangling these interactions will provide a ripe area for future studies to understand how to best harness beneficial endophytes for agriculture.

## Background

Food security is critical to modern global society. However, problems such as soil degradation, climate change, and a growing population will challenge our global food supply in the 21^st^ century [1,2]. Improving crop yield by even small percentages results in massive increases in product and reduces the environmental burden of production. Biostimulants are a class of agricultural inputs based on either living organisms or products derived from them, and the use of biostimulants is touted as a sustainable way to improve yield, reduce soil degradation, and provide other ecological benefits [3].

A common class of biostimulants involves the use of endophytes, microbes that live inside plants’ tissue [4]. Previous studies have identified many ways that endophytes impact their hosts, such as by supplying nutrients (including nitrogen) [5,6,7,8], increasing stress resistance [9,10,11,12] and increasing crop growth and yield [6, 11, 13,14]. Endophytes can stimulate plant growth in a variety of ways, such as by out-competing pathogens [15], producing antimicrobial compounds [16,17,18], synthesizing/increasing phytohormones and secondary metabolites [12,19,20], and mitigating stress [9,10,11,12].

The biostimulant market is projected to be worth 11 billion USD by 2027 [21]. Many such treatments are already commercially available in the form of foliar sprays, soil treatments, and seed treatments. Both startup companies and large corporations like Bayer and Syngenta are heavily investing in biostimulants [21], and trends in the scientific literature indicate growing interest in endophyte growth promotion in the public sector as well [22]. Despite this, it is incredibly challenging to bring a biostimulant from the lab to market [23,24]. Many microbes that appear promising in the lab do not produce reliable effects in the field. Although many factors are probably responsible for this issue, one that has received less attention is how different plant genotypes respond to beneficial endophytes.

Several groups have shown that the effects of beneficial microbes vary across genotypes [25,26,27], but there has been relatively little quantification of this effect and almost no exploration of the underlying mechanisms. For example, an increase in maize yield due to bacteria in the genus *Herbaspirillum* depended on both the endophyte strain and the maize variety, with *Herbaspirillum seropedicae (*ZAE 94) increasing biomass in less than a third of commercial maize genotypes tested [25]. One of the few studies investigating this interaction traced the variation in *Arabidopsis*’ response to growth-promoting rhizobacteria to several candidate genes, including genes involved in plant-growth processes like transporters and metabolism [28]. Understanding the nature and extent of this variation could provide a way to improve the development and use of biostimulants in agriculture.

Maize is one of the most important crops in global agriculture. Over a billion tonnes of maize were produced worldwide in 2019 [29]. Although not investigated in detail, previous research has shown that the same endophyte impacts different maize inbreds to different degrees, and these differences are likely due to differences in the host’s genetics [10, 25,26,27,30,31,32]. Despite the number of studies that mention such genotype-by-microbe interactions, few if any of them quantify the size of the effect, so it is difficult to determine just how important of a factor this variation is. Understanding and effectively utilizing these interactions would be a step towards increasing yield in a sustainable fashion.

In this study we aimed to quantify the effect maize genotype has on host-endophyte interaction. In a series of three experiments, we inoculated diverse maize lines with one of three different endophytes (*Herbaspirillum seropedicae, Burkholderia WP9*, and *Serendipita bescii*) and quantified the resulting changes in phenotype.

*Herbaspirillum seropedicae* is a well-studied gram-negative, growth-promoting endophyte that is commonly used to study nitrogen fixation in symbioses with grasses [14, 25,33]. It has a broad host range and can colonize sugarcane, rice, wheat, and maize, where it can act as a biofertilizer [3,5,34]. In addition to fixing nitrogen, *H. seropedicae* can solubilize minerals and produce phytohormones [35]. It has been shown that maize yield, metabolite content, and shoot dry weight depend upon the genotype of both the *Herbaspirillum* inoculant, as well as the genotype of the maize host [25,26,27].

*Burkholderia* is bacterial genus containing well-studied growth promoting endophytes. Different species have been shown to enhance growth, yield, and disease resistance [6], and aid in the uptake of phosphate and nitrogen [6,36]. *Burkholderia* WP9 was isolated from black cottonwood and has nitrogen fixing abilities [37].

*Serendipita bescii* is a Basidiomycota fungus, and fungi in this genus are known to associate with many different plant species as an endomycorrhizae [38]. Originally recognized as orchid mutualists, *Serendipita* fungi also promote growth in a number of different plants, including switchgrass [39]. It is hypothesized that when serendipitoid fungi colonize root systems, they break down organic manner in the soil and make these nutrients available to the plant [40].

For each microbe, we determined its effect on seedling phenotypes of diverse maize varieties and quantified the effect of maize genetics, microbes, and their interaction. These results provide a firm estimate of the degree to which different maize genotypes respond to beneficial endophytes, and how genetic variation among these lines modulates that response.

## Methods

### Experimental Design

Maize genotypes were selected from among the Goodman-Buckler diversity panel [41], with most also being founders of the maize Nested Association Mapping population [42] (Table 3). For space reasons, each experiment was subdivided into a series of “grows” including only a subset of plant genotypes. Within each grow, plants were arranged in a randomized complete block design with 5 reps; in a few cases seedlings died after germination leaving that genotype with four reps.

### Seed Sterilization and Plant Growth

Seeds were surface sterilized for five minutes using 50mL sterile H_2_O, 50mL of bleach(Clorox), and three drops of Tween 20(VWR). Seeds were rinsed five times with 100mL of sterile water, then immersed in a 60° C water bath for 15 minutes to kill existing endophtytes [43]. Seeds (in water) were then allowed to cool and imbibe for 1 hour before placing 10 seeds equidistant from each other in an autoclaved magenta box with 15 ml nutrient agar (1x Hoagland solution [bioWorld 30630038-5] + 15g/L of agar [Caisson Labs])). The box was then parafilmed shut, and the seeds were allowed to germinate for seven days. After 7 days, seedlings were moved to the greenhouse and planted 4 cm deep in 2.37 liter pots filled with autoclaved Professional Growing Mix Fafard 3B/Metro-Mix 830 (Sungro Horticulture) and inoculated as described below. Pots were watered three times a week, and plants were grown for an additional 21 days before they were harvested.

### Bacterial growth and inoculation

#### Experiment 1

*H. seropediceae* (ATTC 35892) was grown from a single colony in nutrient broth at 24 °C for 48 hours to an OD of ∼0.8. Germinated seeds were placed into the autoclaved soil and inoculated with 2mL of culture or sterile nutrient broth (control) before being covered by soil. No additional water was applied for 2 days to allow for colonization.

#### Experiment 2

*Burkholderia* WP9 (Sharon Doty, University of Washington) was grown from a single colony in nutrient broth at 24 °C for 48 hours to an OD of ∼0.8. Autoclaved soil was inoculated with 200mL of culture/kg of soil, or with sterile nutrient broth for controls, before being placed into individual pots and germinated seedlings planted as above.

#### Experiment 3

*Serendipita bescii* (Kelly Craven, Noble Research Institute) was pre-inoculated onto clay bentonite particles [38,39] by collaborators at the Noble Research Institute. Based on their recommendation, soil was placed into pots a week before sowing and thoroughly washed with water 5 times to leech minerals and nutrients from the media. 100g of clay particles (inoculated or control) were placed in a depression in the soil, with the germinated seed then placed on top and covered.

After 21 days, the above- and below-ground portions of each plant were separated with a sterile razor blade and frozen at -80° C for further phenotyping.

### PCR confirmation of colonization

For each experiment, endophyte colonization was confirmed by PCR. About 0.5g of washed root was collected using a sterile razor blade ∼8cm from the base of the root. Samples were placed into a 2mL microtube with a sterile metal ball (Daisy BBs) and placed into a GenoGrinder 2010 at 1400 RPMs for 5 minutes. DNA was extracted with a Quick-DNA Fungal/Bacterial kit (Zymo) and DNA quality and concentration checked via Infinite M200 Pro (TECAN). PCR was run on each sample using the specific endophyte primers (Additional File 1) and run on a 1% agarose gel to confirm colonization. Non-inoculated plants served as controls for greenhouse contamination. PCR program was as follows: 30s at 95°C, followed by 30 cycles of 15s at 95°C, 60s at 59°C, 30s at 68°C, ending with 5m at 68°C and hold at 4°C.

### Phenotyping Methods

#### Plant Height

was measured from the soil line to the tip of the longest/tallest leaf when held upright, and was recorded every week. This method is thus is a combination of plant height and leaf length, and was used to control for different leaf angles.

#### Chlorophyll

Quantum Yield was measured using a Flouropen FP 100 (Photon Systems Instruments). Measurements were taken from halfway up the most mature leaf on the plant. Three measurements were taken at the same location and averaged.

#### Leaf Area

The newest mature leaf was gently removed at the collar. Leaves were laid flat and pinned to a white surface next to a 1”-square size marker and a paper with sample identifying information (name & date). Images of each leaf were quantified with EasyLeafArea [44], with the following batch parameters: Leaf minimum Green RGB value 15, Leaf Green Ratio (G/R) 1.06, Leaf Green Ratio (G/B) 1.08, Scale Minimum Red RGB value 50, Scale Red Ratio 1.96, Scale area (cm^2^) = 6.5 cm^2^.

#### Shoot Mass

The entire aboveground portion of the plant, including the leaf removed to measure leaf area, was dried in a forced-air oven at 37.7 ° C for 48 hours before being weighed on a precision balance (VWR 164AC).

#### Roots

Frozen roots were removed from the freezer and washed with warm sterile water while gently rubbing to remove as much soil as possible without damaging the root system. Root length was measured from the base to the end of the longest root. Root volume was measured by placing the washed & dried roots into a 20-ml graduated cylinder half-filled with water and recording the displacement volume as the roots were submerged. Dry root mass was measured on a precision balance (VWR 164AC) after air-drying samples for 48 hours at 37.7 °C in a forced-air drier.

#### qPCR for fungal biomass

For Experiment 3 only, relative fungal biomass was estimated using the 2^ΔΔCt^ method [45] to compare the amount of fungal ITS3 to maize CDK DNA in each sample. Extracted maize root DNA (the same used to confirm colonization, above) was diluted to 12 ng/uL using an Infinite M200 Pro (TECAN). qPCR was performed using primers specific for the *Serendipita* ITS3 gene [39] or the maize CDK (cycline-dependent kinase housekeeping gene) [46] (Table S1). Reactions were performed using SYBR Green I Master Mix (Roche) and the manufacturer’s recommended protocol (pre-incubation at 95°C for 5m, 45 cycles of amplification for 10s at 95°C, 49.6°C (ITS) or 59.3°C (CDK) for 18s, and 30s at 72°C). A single melting curve was performed, with 8 acquisitions/°C. Reactions were run on a Roche LightCycler 480, with two technical replicates for each sample. 2^ΔΔCt^ values were generated from threshold crossing (Ct) values, and then log transformed to visualize *S. bescii* colonization for each maize genotype in R.

### Statistics

All statistics were run in R [47]. Due to space constraints, groups of genotypes (“grows”) had to be planted separately throughout each experiment; since these grows were completely confounded with the plant genotype, they were not included in subsequent analyses. ANOVA was performed by fitting a linear model of Phenotype ∼ Genotype + Condition + Genotype:Condition, where “Condition” represents either inoculated or control. The resulting ANOVA table was used to calculate the fraction of total variation contributed by each component and its statistical significance. The Genome x Genome interaction represents the fraction of variation explained after accounting for the main effects of maize genotype and the general effect of inoculation. GxG is calculated as V_GC_ / (V_GC_+V_e_), where V_GC_ is the variation due to genotype-by-condition interaction, and V_e_ is the residual (error) variance. Significant growth differences were determined using a Welch’s two-sided t-test between inoculated and control plants of the same genotype. Levene’s test was used to check for homogeny of variance for each phenotype.

## Results

We tested three separate endophytes for growth promoting abilities in maize inbred lines in three separate greenhouse experiments. Due to space constraints, experiments had to be subdivided into Grows, which contained all of the replicates for a genotype. The experimental design was kept identical for grows within the same experiment, with planting date being the only difference. All tested phenotypes are listed in Table 1. In these experiments, “genome-by-genome interaction” (“GxG”) refers to how much variation was due to the specific combination of maize variety and endophyte inoculation. Functionally, GxG represents the differences in how maize varieties respond to an endophyte, so that the larger GxG is the stronger the differences are among maize varieties.

**Table 1:** Maize germplasm

| Maize Genotype | GRIN Accession | Experiment(s) Used |
| --- | --- | --- |
| **A635** | PI 693329 | 1, 2, 3 |
| **B73** | PI 5504073 | 1, 3 |
| **B97** | PI 564682 | 1, 3 |
| **CML 52** | PI 595561 | 1, 3 |
| **CML 103** | Ames 27081 | 1, 3 |
| **CML 228** | Ames 27081 | 1, 2 |
| **CML 333** | Ames 27101 | 3 |
| **HP301** | PI 587131 | 3 |
| **KI3** | Ames 27123 | 1, 3 |
| **KI11** | Ames 27124 | 2, 3 |
| **MO17** | PI 558532 | 1 |
| **MS71** | PI 587137 | 1, 2, 3 |
| **NC350** | Ames 27171 | 1, 2, 3 |
| **P39** | Ames 28186 | 2, 3 |
| **TX303** | Ames 19327 | 1, 3 |

### Experiment 1

Experiment 1 used the gram-negative bacterium *Herbaspirillum seropediceae* Z67, with liquid culture applied directly to germinated seeds. Phenotypic variation was largely due to the plant genotype (Figure 1), as was expected given the high genetic variation in maize. Inoculation with *H. seropediceae* was not a significant source of variation for any phenotype, and GxG interaction was only statistically significant for root volume (p=.05; Figure 1). Root volume had the highest GxG for the entire study (0.243, after accounting for the main effects of genotype and endophyte; Table 1). When looking at individual genotypes instead of the experiment as a whole,*H. seropediceae* increased chlorophyll content, plant height, and root volume (Table 3), though only for 1 maize genotype in each case. *Herbaspirillum* increased growth, though generally only for a couple of maize genotypes. There was also an interesting trend where *H. seropediceae* increased root length in A635 but decreased it in CML52, though it did not reach statistical significance.

**Figure 1.**
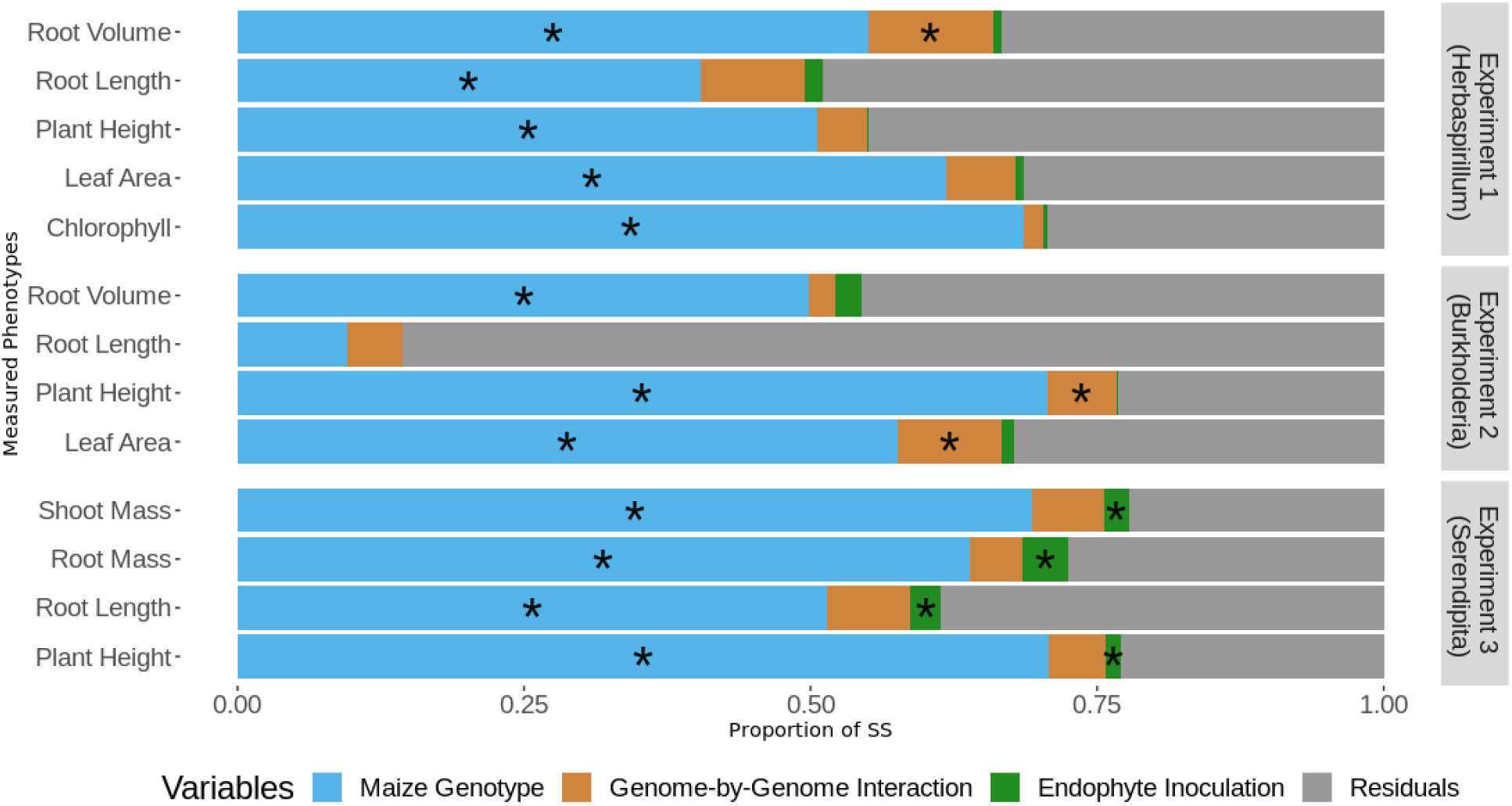
ANOVA analysis.Standard ANOVA was used to break phenotypic variation into the effects of maize genotype, endophyte inoculation, their interaction, and residual error. Asterisks denote significant effects (p<0.05). Plant genotype is almost always significant, while only *Serendipita* (Experiment 3) shows a consistent effect of the endophyte. Levene’s test was used to check for equality of variances (one of the assumptions of ANOVA), and only Shoot Mass in Experiment 3 failed the test, so its results should be taken with caution.

### Experiment 2

Experiment 2 used the gram-negative bacteria *Burkholderia* sp. WP9, with liquid culture applied directly to bulk soil immediately before adding seeds. Again, plant genotype was statistically significant for most measured phenotypes while endophyte inoculation by itself was not significant for any (Figure 1). Plant-endophyte interaction was statistically significant for both Plant Height and Leaf Area, with GxG interaction scores of ∼0.2 (Table 1). Although *Burkholderia WP9* reportedly boosts maize growth (Sharon Doty, personal communication), we saw no statistically significant growth promotion for any individual phenotypes in any lines that were tested (Table 2). The fact that GxG interaction overall is significant while no individual genotypes are is probably due to the greater statistical power when looking at the experiment as a whole.

**Table 2.** Genome x Genome interaction calculations. GxG were calculated as the fraction of phenotypic variance due to Genotype:Inoculation interaction after accounting for the main effects of maize genotype and inoculation.

| Endophyte | Trait | GxG Interaction (0-1) |
| --- | --- | --- |
| <u>Experiment 1</u> |  |  |
| *Herbaspirillum* | Chlorophyll | .057 |
| *Herbaspirillum* | Plant Height | 0.091 |
| *Herbaspirillum* | Leaf Area | 0.155 |
| *Herbaspirillum* | Root Length | 0.157 |
| *Herbaspirillum* | Root Volume | 0.243 |
| <u>Experiment 2</u> |  |  |
| *Burkholderia* | Plant Height | 0.218 |
| *Burkholderia* | Leaf Area | 0.232 |
| *Burkholderia* | Root Length | 0.071 |
| *Burkholderia* | Root Volume | 0.027 |
| <u>Experiment 3</u> |  |  |
| *Serendipita* | Plant Height | 0.178 |
| *Serendipita* | Root Length | 0.157 |
| *Serendipita* | Root Mass | 0.144 |
| *Serendipita* | Shoot Mass | 0.222 |

**Table 3.**
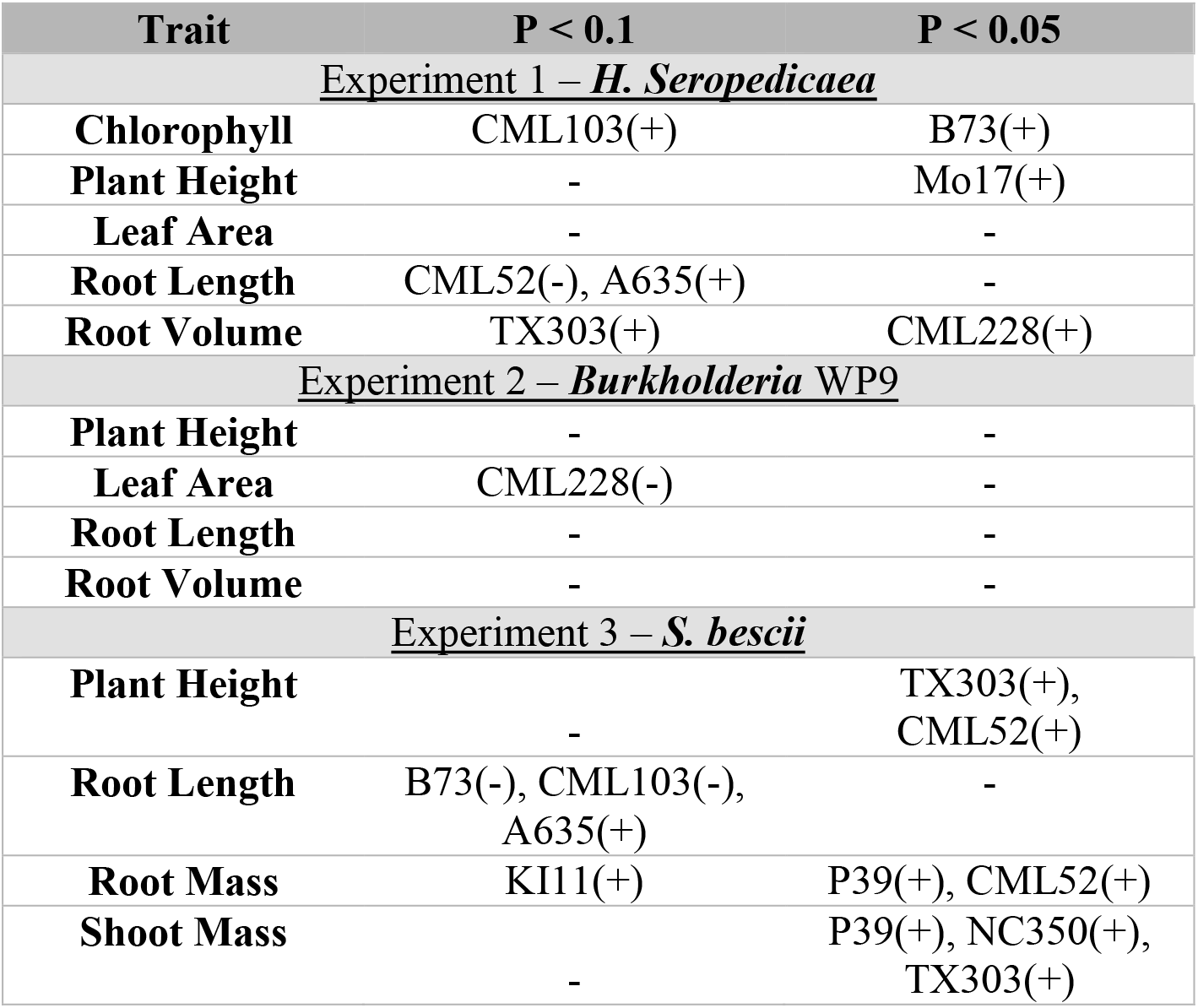
Genotypes that have significant growth promotion for a measured phenotype. Significance was determined by a two-sided t-test of control versus inoculated plants (see Methods). The direction of growth (increase or decrease) is indicated by a “+” or “–” sign after each genotype.

| Trait | P < 0.1 | P < 0.05 |
| --- | --- | --- |
| <u>Experiment 1 – *H. Seropedicaea*</u> |  |  |
| **Chlorophyll** | CML103(+) | B73(+) |
| **Plant Height** | - | Mo17(+) |
| **Leaf Area** | - | - |
| **Root Length** | CML52(-), A635(+) | - |
| **Root Volume** | TX303(+) | CML228(+) |
| <u>Experiment 2 – *Burkholderia* WP9</u> |  |  |
| **Plant Height** | - | - |
| **Leaf Area** | CML228(-) | - |
| **Root Length** | - | - |
| **Root Volume** | - | - |
| <u>Experiment 3 – *S. bescii*</u> |  |  |
| **Plant Height** | - | TX303(+),<br>CML52(+) |
| **Root Length** | B73(-), CML103(-),<br>A635(+) | - |
| **Root Mass** | KI11(+) | P39(+), CML52(+) |
| **Shoot Mass** | - | P39(+), NC350(+),<br>TX303(+) |

### Experiment 3

Experiment 3 used the Basidiomycota fungus *Serendipita bescii*, which was pre-inoculated on sterile clay particles that were placed directly under the germinated seed in the soil. *S. bescii* showed the strongest growth-promoting effects of the three endophytes, with a significant main effect of inoculation for all four measured phenotypes (Figure 1). *S. bescii* increased growth in a number of genotypes (Table 2; Figure 2), though GxG interaction was not significant in the experiment overall (Figure 1), possibly due to the high variance within some of the lines. A particularly interesting contrast in this experiment involves the maize lines CML52, NC350, P39, and TX303 (Table 2). Inoculation with *S. bescii* increased only the above-ground biomass of NC350 and TX303, only the belowground biomass of CML52, and both traits for P39 (Figure 2).

**Figure 2.**
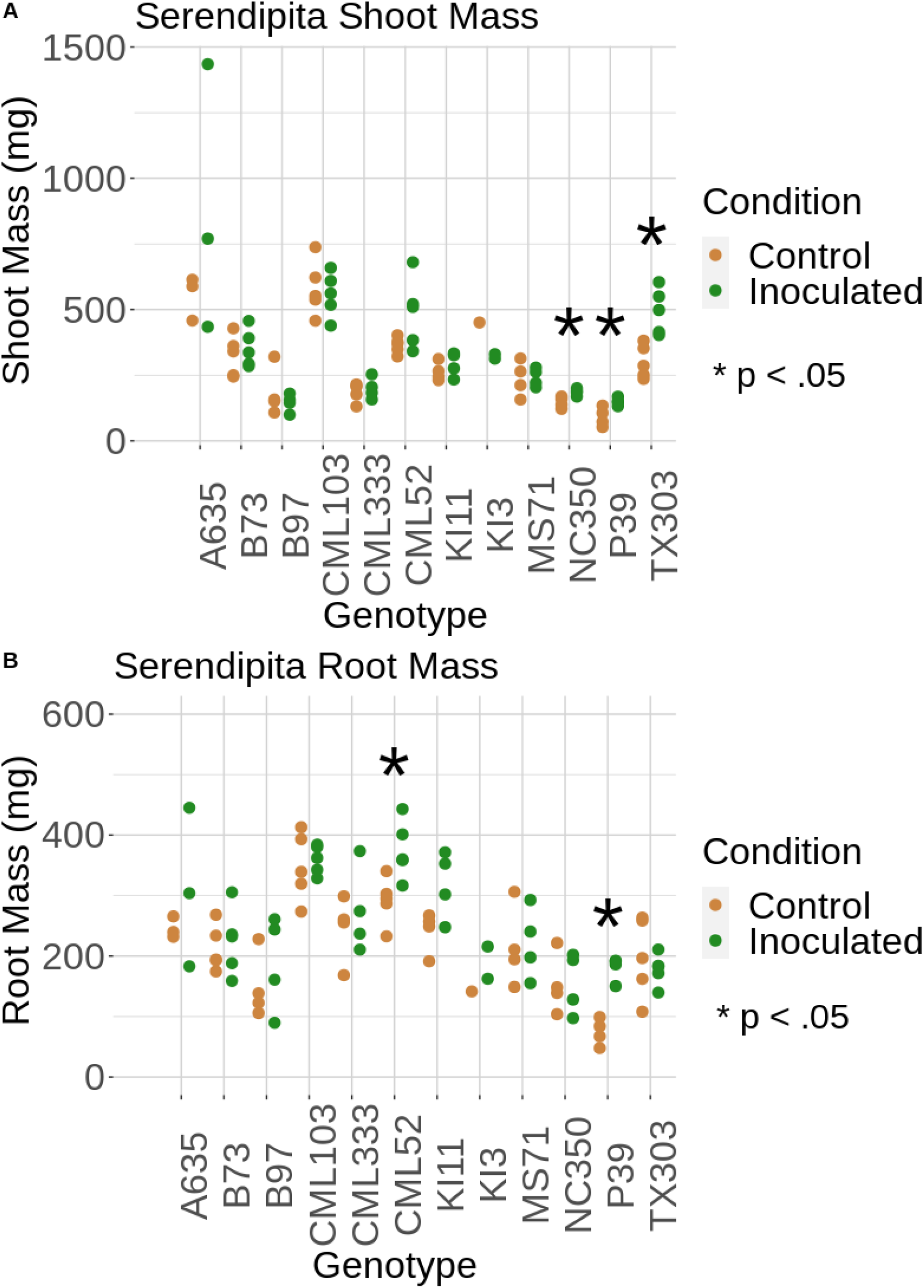
*Serendipita* growth promotion. Effect of inoculation of *S. bescii* on different maize gentoypes for (A) shoot and (B) root biomass. Asterisks denote significant growth promotion. Several genotypes show growth promotion in only one of the two compartments; only P39 shows promotion in both.

These differences in phenotype response could be due to differences in endophyte colonization, as has been shown for other grasses [48, 49]. We quantified the colonization of *Serendipita* using qPCR of the *Serendipita* fungal ITS3 [39] normalized against the maize housekeeping gene CDK [46] (Figure 3). Higher levels of *Serendipita* colonization coincided with increased growth in the greenhouse. Maize lines P39, NC350, TX303, and CML52 had high *Serendipita* in their respective grows, and showed growth promotion for at least one phenotype with a p-value < 0.05 (Table 2); two other maize genotypes (A635 and KI11) also had high *S. bescii* abundances but less significant growth promotion (p < 0.1; Table 2). Of the remaining lines, MS71 and CML333 showed no growth promotion, while CML103 and B73 had a nonsignificant trend (p ∼ 0.07) toward growth inhibition.

**Figure 3.**
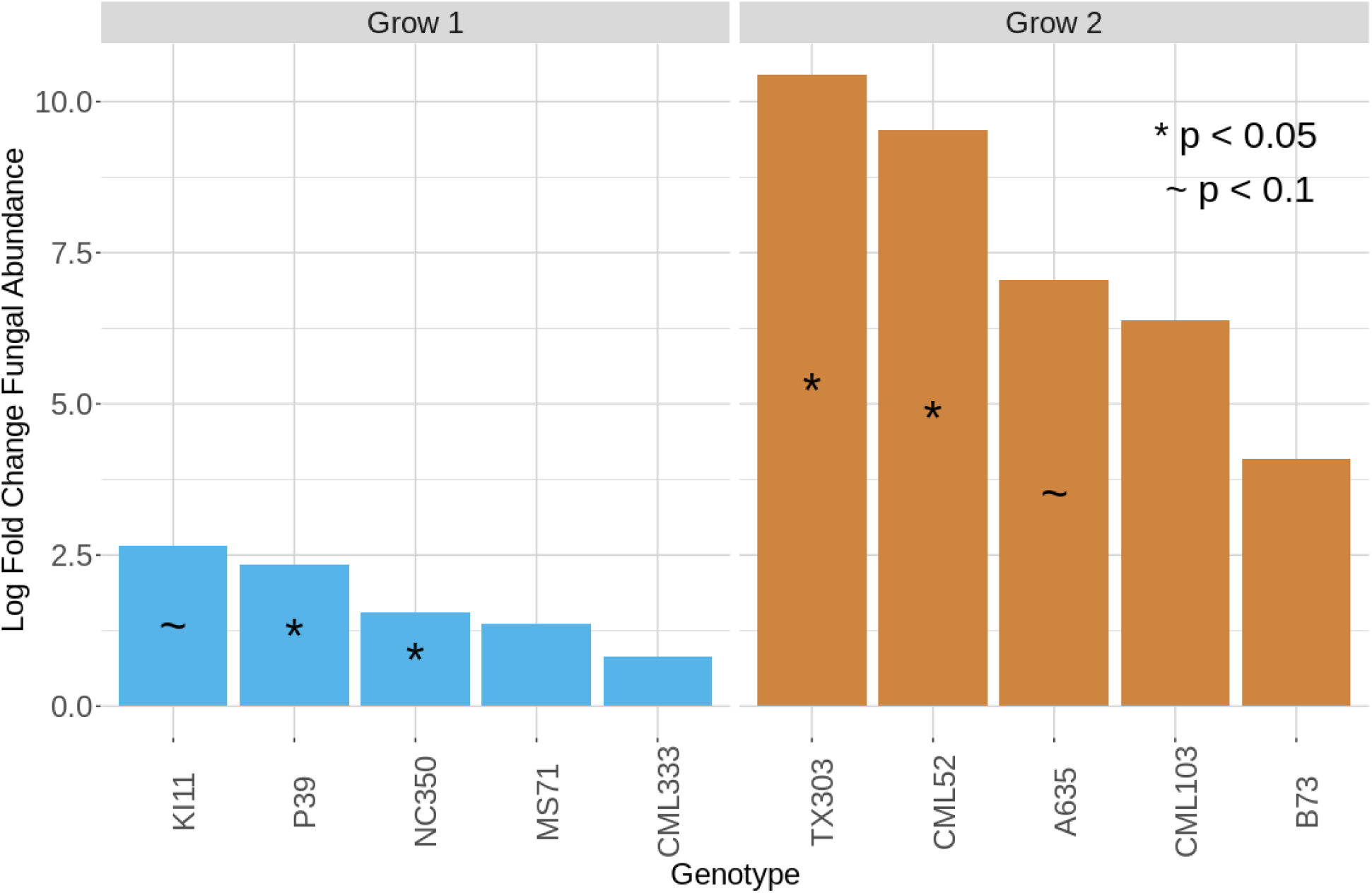
Differences in *Serendipita* Abundance by Genotype. Log fold-change differences in *S. bescii* colonization between control and inoculated plants. The *S. bescii* ITS3 gene was quantified via qPCR and normalized to the maize CDK housekeeping gene. Since “grows” were planted on separate dates for space reasons, they are kept separate in this analysis. Within each grow, higher levels of colonization coincided with significant growth promotion (marked with ∼ or *). The least-colonized plants in each grow showed either no promotion or trends toward growth inhibition (Table 2).

## Discussion

Our results in this study highlight how three potentially growth-promoting endophytes affect genetically diverse maize. When inoculated with *Herbaspirillum seropedicae* (Experiment 1), we found several maize genotypes showed increased growth in one of several phenotypes (Chlorophyll content, plant height, root length, and root volume, Table 2). Interestingly, each genotype showed an increase in a specific phenotype, and genome-by-genome interaction played a significant role only for root volume (Figure 1). Experiment 2, examining *Burkholderia* WP9, indicated significant GxG interaction for two phenotypes (plant height and leaf area), though an examination of individual varieties did not show statistically significant effects, probably due to the smaller sample size. Since other studies have shown that this isolate promotes growth in both rice [48] and maize (Sharon Doty, personal communication), it may be worth additional investigation. Experiment 2 did show that some maize genotypes were slightly hindered by the endophyte, while some received slight growth promotion, highlighting the importance of genotype interactions. Of the genotypes inoculated with *Serendipita bescii* (Experiment 3), one (P39) showed growth increases both above and below ground (Figures 2 & 3), while three other genotypes (NC350, CML52 and TX303) experienced increases in only one of the two categories. Similarly to *Burkholderia*, inoculation with *Serendipita* decreased the root length of two maize genotypes (B73 and CML103), further emphasizing the potential for unique reactions to endophyte treatment among different genotypes. *Serendipita* was the only endophyte to show a main effect of inoculation across all phenotypes (Figure 1), meaning it was the only one of the three endophytes to have a relatively consistent effect on hosts regardless of genotype.

Not only did *Serendipita* have the most consistent effect on hosts, we also showed that the size of this effect may be related to the plants colonization level (Figure 4). This indicates that stronger growth-promotion phenotypes may be a direct consequence of higher endophyte loads. We attempted to find other examples of this trend in the literature, but there are surprisingly few studies that directly compare endophyte colonization with growth promotion. One exception involves Epichloë fungal endophytes, whose biomass has been shown to correlate with the amount of protective alkaloid compounds in their grass hosts [49,50]. Further testing is needed to confirm if endophyte biomass amounts have a direct influence on growth promotion, and to uncover the mechanisms and genetic diversity behind these interactions in maize.

This data does not provide insight into the molecular mechanisms of genome-by-genome interaction, but interest in this field is quickly growing. Prior work indicates that these interactions could be impacted by a number of metabolites and pathways. For example, diverse maize genotypes respond differently to microbe-associated molecular patterns (MAMPs), and as a consequence show significant differences in reactive oxygen species, nitric oxide production, and defense gene expression [51,52]. It has been suggested that *Herbaspirillum* may regulate the ROS pathway differently in diverse maize roots [26]. Many endophytes produce phytohormones [4,12] and volatile organic compounds [20,35], which directly impact growth promotion and stress resistance. Future studies may be able to shed light on how host genetic diversity impacts these molecular interactions.

A chief goal of this research was to quantify “Genome-by-Genome interaction” in this system, meaning changes in phenotype that depend on both the maize genotype and the specific endophyte. The GxG interactions we identified explain relatively little phenotypic variance, even after factoring out the effect of maize genotype (Table 1). The strongest GxG effects explain ∼20-25% of residual variance after accounting for genotype, although most are half this or less. Looked at another way, GxG interaction generally explained only 1/10th of the variance that maize genotype did, meaning that the effect was 10 times weaker than the effect of plant genetics. This small effect size may make it challenging to attempt to dissect the underlying genetic components that affect plant-microbe interactions, a conclusion shared by a recent review of GWA studies [53]. One should keep in mind, however, that the maize varieties in this study were specifically chosen for their high diversity. Diversity in elite breeding programs is generally much lower, so endophyte inoculation may have relatively larger effects within elite material.

These data demonstrates that maize genotype does have a significant impact on endophyte-mediated growth promotion. In addition, all three endophytes showed indications of decreased growth in at least one maize line, but none of these lines were the same. On the other hand, of the nine maize genotypes tested with both *H. seropedicae* and *S. bescii*, only one (TX303) showed statistically significant growth promotion for both endophytes. Interestingly, this promotion was belowground (Root Volume) for *H. seropedicae* and above ground (Plant Height) for *S. bescii*. This may indicate that these endophytes are activating different pathways or altering the expression of different genes.

## Conclusion

Our interactions with maize producers in the US indicates that they are both interested in biologicals and concerned about their efficacy. Our findings show how different maize inbreds react to the same beneficial microbe. These findings provide insight into the range of responses between plants and microbes in an agricultural setting, and are especially important for groups developing new bioinoculates and biofertilizers. The high variability across genotypes may partly explain why many beneficial microbes reported in the literature fail to translate to field production. (This is apart from logistical factors such as scalability, shelf life, ease of use, and compatibility with existing formulation [23, 24], all of which also play a role.) These results imply that new microbial-based products for agriculture should be screened against diverse genotypes early in the process so as to weed out microbes with variable effects. Conversely, it may be possible to eventually tailor microbes to interact well with specific genotypes and deliver them as a specific plant-microbe combination (using seed treatments, for example). Further studies will show how these interactions occur and how we can best harness them to improve global agriculture.

## Supporting information

CombinedRaw_qPCR_clean

Final_Burk_Data

Final_Herb_Data

Final_Sbecii_Data

## Declarations

### Ethics approval and consent to participate

Not applicable

### Consent for publication

Not applicable

### Availability of data and materials

The datasets generated and analyzed during the current study, along with all code are available at https://github.com/wallacelab/paper-schultz-endophytes-2021

### Competing interests

The authors declare that they have no competing interests.

### Funding

Funding for this experiment was provided by the University of Georgia and the Foundation for Food and Agriculture Research (FFAR).

### Authors’ contributions

J.W. conceived and supervised the work. C.S. performed the experiments and data analysis. K.B. assisted with Experiment 2. J.W. and C.S. co-wrote the manuscript. All authors reviewed the results and approved the final version of the manuscript.

## Acknowledgments

We would like to thank Dr. Sharon Doty (University of Washington) for supplying the *Burkholderia WP9* for this experiment, and Drs. Kelly Craven and Prasun Ray (Noble Research Institute) for supplying the *Serendipita bescii* inoculum.

## Additional Files

Additional File 1

Additional_File_1.csv

Equipment List and Endophyte Primers

This file contains information on reagents, equipment, and primers used in this study.

## Notes

https://github.com/wallacelab/paper-schultz-endophytes-2021

### Competing Interest Statement

The authors have declared no competing interest.

